# Post-weaning social isolation increases reward-seeking behavior in a sex-specific manner in mice

**DOI:** 10.1101/2021.04.20.440665

**Authors:** Michael Noback, James C. Barrow, Gregory V. Carr

## Abstract

Social isolation is a growing concern in public health. Although isolation at any age is harmful, previous studies have shown that isolation during adolescence, correlating with critical periods of brain development, can impair cognitive function and increase the risk for psychiatric illness later in life. In this study, we utilized a mouse model of adolescent social isolation (SI) and compared performance of isolated and group-housed mice on touchscreen-based continuous performance test (CPT) and fixed ratio/progressive ratio (FR/PR) tasks in adulthood. SI increased sensitivity in the CPT in male mice and had no effect in female mice. The increase in sensitivity was consistent across time bins within the 45-minute testing session and there were no SI effects on reaction times or reward retrieval latencies. A possible confound for performance in the CPT would be SI-induced changes in reward-seeking or motivation for the strawberry milk reward. We next compared the SI mice to their group-housed littermate controls on both FR and PR schedules of reinforcement and found that male SI mice earned significantly more reinforcers on FR schedules of reinforcement and had higher breakpoints on PR schedules compared to their group-housed littermates. SI had no effect on FR or PR performance in female mice. These data indicate that SI during adolescence has striking, sex-specific effects on reward-seeking behavior in adult mice and may provide a useful behavioral model for studying the link between SI and risk for neuropsychiatric disorders.

## 1. Introduction

Social isolation (SI) is a stressor with significant acute and chronic effects, including increased risk for developing psychiatric disorders. Specifically, there is a clear link between SI and anxiety, depression, substance use disorder, and schizophrenia [1]. Despite the confirmation of SI as a risk factor, the biological link between SI and psychiatric disorders is unknown.

The relevance of SI as a source of environmental stress has increased dramatically over the past year due to the COVID-19 pandemic. It is too early to definitively assess the societal and economic impacts of the increase in SI due to social distancing measures, but its role in public health is already being closely monitored [2–4]. A deeper understanding of the biological effects of SI will be critical to address the long-term effects of COVID-19-related SI.

There is an extensive literature on the effects of SI on older people (Reviewed in [5]), but recent studies have identified increases in the prevalence of SI and loneliness among children and adolescents [6]. SI in younger people is particularly important for disorders with significant neurodevelopmental components, including schizophrenia and substance use disorders [7]. Given the increased plasticity of the brain during childhood and adolescence [8], SI may have different or greater effects compared to SI in adults. SI has also been studied in animal models, and both chronic and acute isolation have been associated with anxiety- and depression-like behavior in rodents [9]. SI during postnatal days 21-35, roughly correlating developmentally to ages 11 to 18 in humans, [8] produced deficits in myelination in the prelimbic and infralimbic cortices through an IL-6-dependent mechanism [10,11]. Additionally, SI during this period also enhances the activity of interneurons in the prelimbic and infralimbic cortices [12].

It is not clear why isolation during postnatal days 21-35 produces such drastic effects on the structure and function of the brain. Interestingly, postnatal days 21-35 is a period of significant amounts of social play behavior in mice and isolation during this period can lead to deficits in social interactions in adulthood [13,14]. Moreover, the social behavior deficits are directly related to abnormal interneuron function in the prefrontal cortex [14]

We previously reported that SI during the 21-35 day period is associated with increased levels of ΔFosB, a transcription factor associated with chronic stress[15]. ΔFosB has also been implicated in studies of reward-seeking behavior[16–18]. Given the developmental trajectory of the cortex and connected reward circuits [19], these studies suggest there may be a link between SI, ΔFosB, and cognitive function, particularly reward-seeking behavior.

Touchscreen-based tasks provide a flexible, sensitive, and translational platform for investigating cognitive function [20]. Sustained attention is a cognitive domain disrupted in neuropsychiatric disorders [21,22] The continuous performance test (CPT) is a widely-used and translatable behavioral assay sensitive to changes in sustained attention [23]. Fixed ratio and progressive ratio tasks can similarly detect changes in motivation and reward-related behavior [18]. Here, in a mouse model of adolescent social isolation, we used a cognitive testing battery consisting of CPT and PR assays to interrogate the effects of adolescent SI on attention and motivation in adulthood.

We found that adolescent SI significantly affects performance in CPT, PR, and FR assays. Male SI mice demonstrated higher sensitivity in the CPT, which is usually indicative of improved attention. However, male SI mice also have higher breakpoints and total responses in the PR and FR tasks, indicating increased reward-seeking behavior. Together, these results suggest that improved performance in the CPT in SI mice may result from increased reward-seeking and motivation for highly palatable food.

## 2. Material and methods

### 2.1 Mice

Breeding pairs of C57BL/6J mice were purchased at 6-7 weeks old from The Jackson Laboratory (Bar Harbor, ME, USA). Only one litter from each breeding pair was used in these experiments. Pups were raised according to an isolation protocol adapted from one used by Makinodan and colleagues [10]. Pups were weaned at postnatal day 21 (P21) and housed either in isolation or in a group of three same sex littermates. Isolated mice were rehoused with another same sex isolated littermate at P35 for the duration of the experiment (Figure 1a).

**Figure 1.**
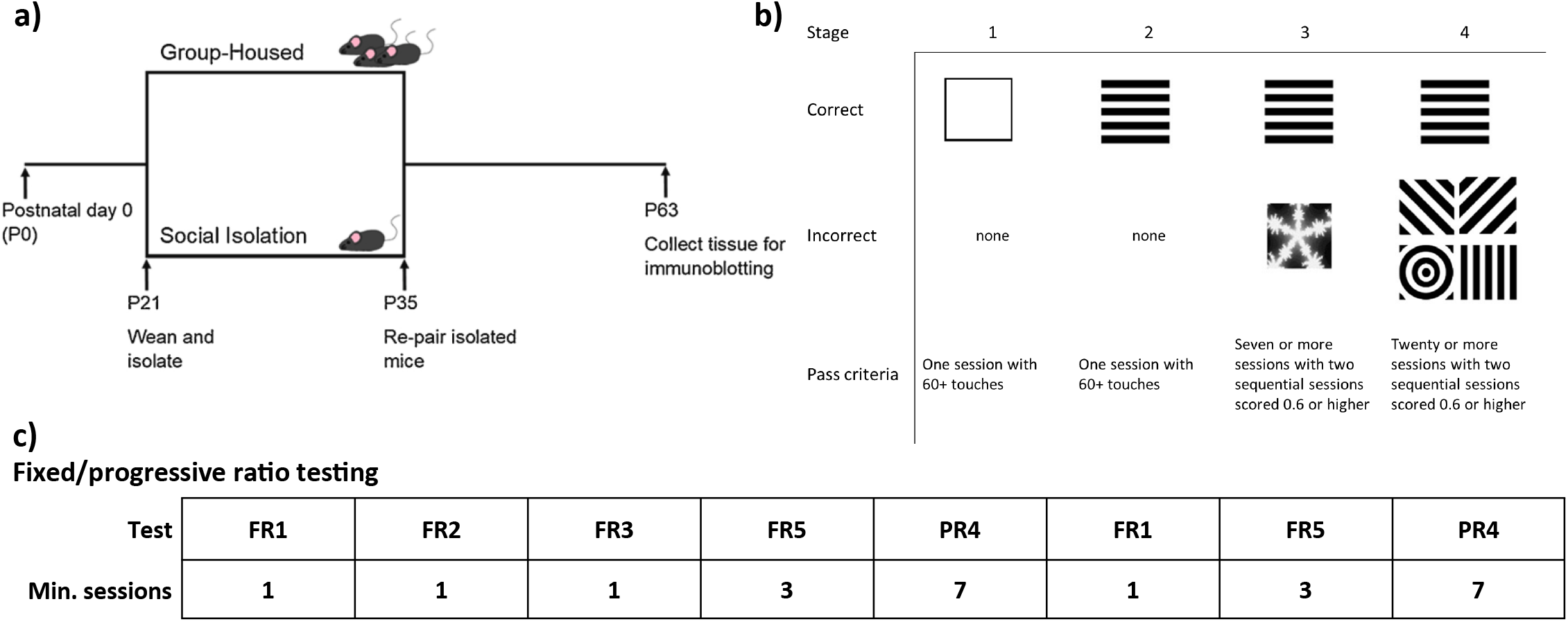
Isolation rearing schematic and behavioral testing battery. a) Isolation rearing. Mice are weaned at postnatal day 21 and assigned to either group housing or isolation. At day 35, isolated mice are rehoused with another isolated littermate. Testing began at day 63. b) Continuous performance test (CPT) progression and scoring. c) Motivation test progression. Session amounts shown are number of passing scores required before moving onto the next stage.

### 2.2 Handling, food restriction, and habituation

At P63, mice were placed on food restriction and briefly handled for three days. Food amounts were set to keep the animals at no less than 85% of their free feeding weight. During the initial handling days, mice were also introduced to a Bussey-Saksida testing chamber (Lafayette Instrument, Lafayette, IN, USA) for habituation. Mice were placed in the chamber, and 1 mL of reward (strawberry Nesquik, Nestle) was placed in the feeding tray. The habituation schedule consisted of a 30-minute session where the screen was responsive to touch, but touches were not rewarded. Mice were considered habituated when they had undergone at least three habituation session and had fully consumed the reward during at least one session.

### 2.3 Continuous performance test (CPT)

CPT training consists of four stages (Figure 1b, adapted from [23]).

In stage 1, mice were trained to touch a visual stimulus (white square). The square was displayed (stimulus duration) for 10 seconds. The stimulus duration was followed by a 0.5 second limited hold (LH) period during which the screen was blank, but a touch would still yield a reward. Upon interacting with the stimulus, a one-second 3 kHz tone would sound, a small amount of reward would be dispensed, and the reward tray would be illuminated. Head entry into the reward tray was detected by an infrared (IR) beam, and upon head entry the intertrial interval (ITI) of 2 seconds would begin. If the mouse did not interact with the stimulus, the ITI would begin immediately following the LH period. The next trial would begin immediately following the ITI. The criterion for advancement to stage 2 was obtaining 60 rewards within a single 45-minute session.

In stage 2, the white square pattern was replaced with either horizontal or vertical bars – the mouse’s assigned positive stimulus, or S+. The S+ was counterbalanced within groups. The SD was reduced from 10 seconds to 2 seconds, and the LH was increased from 0.5 seconds to 2.5 seconds. The criterion for advancement was the same as stage 1.

In stage 3, a negative stimulus (S-; snowflake pattern) was added. Each trial had a 50% chance of being either an S+ or S- trial. SD and LH were identical to stage 2, but ITI was increased to 5 seconds. Interacting with S- during the SD or LH would not yield a reward and would start the ITI. The criteria for advancement to stage 4 were a minimum of seven sessions, during which at least two consecutive sessions had a discrimination index (d’) score of 0.6 or higher. A discrimination index is a measurement derived from signal detection theory [24]that is used to distinguish meaningful response to stimulus from noise. The discrimination index was calculated as follows:

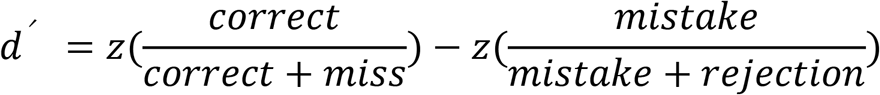

In stage 4, the snowflake S-pattern from stage 3 was replaced with four new S- patterns. The S+ had a 30% chance of appearing, and the remaining 70% was split among the four S- images. Interacting with S- in this stage triggered correction trials, which consisted only of S- trials until the mouse withheld response. Parameters and scoring criteria were identical to stage 3, but animals were tested for twenty sessions.

### 2.4 Fixed ratio (FR)/progressive ratio (PR) testing

FR/PR testing consisted of several stages of varying difficulty (Figure 1c). Fixed ratio 1 (FR1) required a mouse to touch a white square, similar to CPT stage 1 without the stimulus duration timer. The session ended after 45 minutes, or when a mouse obtained 30 rewards within one session. After a mouse obtained 30 rewards within a single session, they progressed onto the next stage. FR2 and FR3 followed the same rules as FR1, but with the respective number of touches per reward. FR5 consisted of multiple sessions in which the mouse had to touch the square five times to obtain a reward. The same timing and cutoff rules from the previous stages applied, but mice were required to obtain 30 rewards during each of at least three sessions before advancing to progressive ratio testing. Progressive ratio 4 (PR4) followed the same parameters as the previous FR stages, but the number of touches required for a reward increased by four each time a reward was obtained. The initial PR4 phase consisted of seven sessions, and during each session the break point of each mouse was recorded. The break point was defined as the highest number of touches committed within one session to obtain a reward.

Following the initial PR4 phase, mice were again tested on FR1, but the 30-reward cap was removed. Three similar sessions of FR5 followed. The amount of rewards obtained per session was recorded. After the free-feeding sessions, mice underwent another seven-session PR4 phase.

## 3. Data analysis

### 3.1 CPT

Discrimination indices from the last ten sessions of stage 4 were used in analysis. ANOVAs were used to analyze the effects of sex and housing condition on CPT performance.

### 3.2 FR/PR

Break points from the fourteen total PR4 sessions were used in analysis. T tests were used to compare group-housed males to isolated males. The numbers of rewards collected during the unlimited FR1 session between PR4 blocks were similarly compared.

## 4. Results

### 4.1 SI did not affect progression through the early stages of the CPT

Stage 1 of the CPT trained mice on the basic mechanism of the test chamber. Neither housing nor sex affected acquisition of test rules in this stage (Housing: F_1,23_ = 0.5490, p = 0.4662; Sex: F_1,23_ = 0.5957, p = 0.4481, Table 1). In stage 2, each mouse tested reached criterion in a single session. In stage 3, neither housing nor sex affected the number of sessions required to reach criterion (Housing: F_1,23_ = 0.0820, p = 0.7772; Sex: F_1,23_ = 0.0820, p = 0.7772), or the d’ score upon reaching criterion (Housing: F_1,23_ = 0.9956, p = 0.3288; Sex: F_1,23_ = 0.1988, p = 0.6599).

**Table 1.** Sessions to completion for the first three stages of CPT. Mean sessions required to reach criterion in each of the training stages of the continuous performance test for each rearing condition. n = 6 GH males, 7 SI males, 6 GH females, and 8 SI females.

|  | Male |  | Female |  |
| --- | --- | --- | --- | --- |
|  | Group | SI | Group | SI |
| Stage 1 | 4.666667 | 4.857143 | 4.833333 | 3.5 |
| Stage 2 | 1 | 1 | 1 | 1 |
| Stage 3 | 9.666667 | 7 | 7 | 9 |

### 4.2 SI improves performance in CPT stage 4 compared to group-housed controls

We found that d’ scores during stage 4 of CPT was increased in SI mice (Figure 2, F_1,23_ = 7.060, p = 0.0141). Sex did not significantly affect performance in CPT (F_1,23_ = 0.8541, p = 0.3650). However, the difference between SI males and group-housed males was greater than that between SI females and group-housed females (Male: p = 0.0668; Female: p = 0.9138).

**Figure 2.**
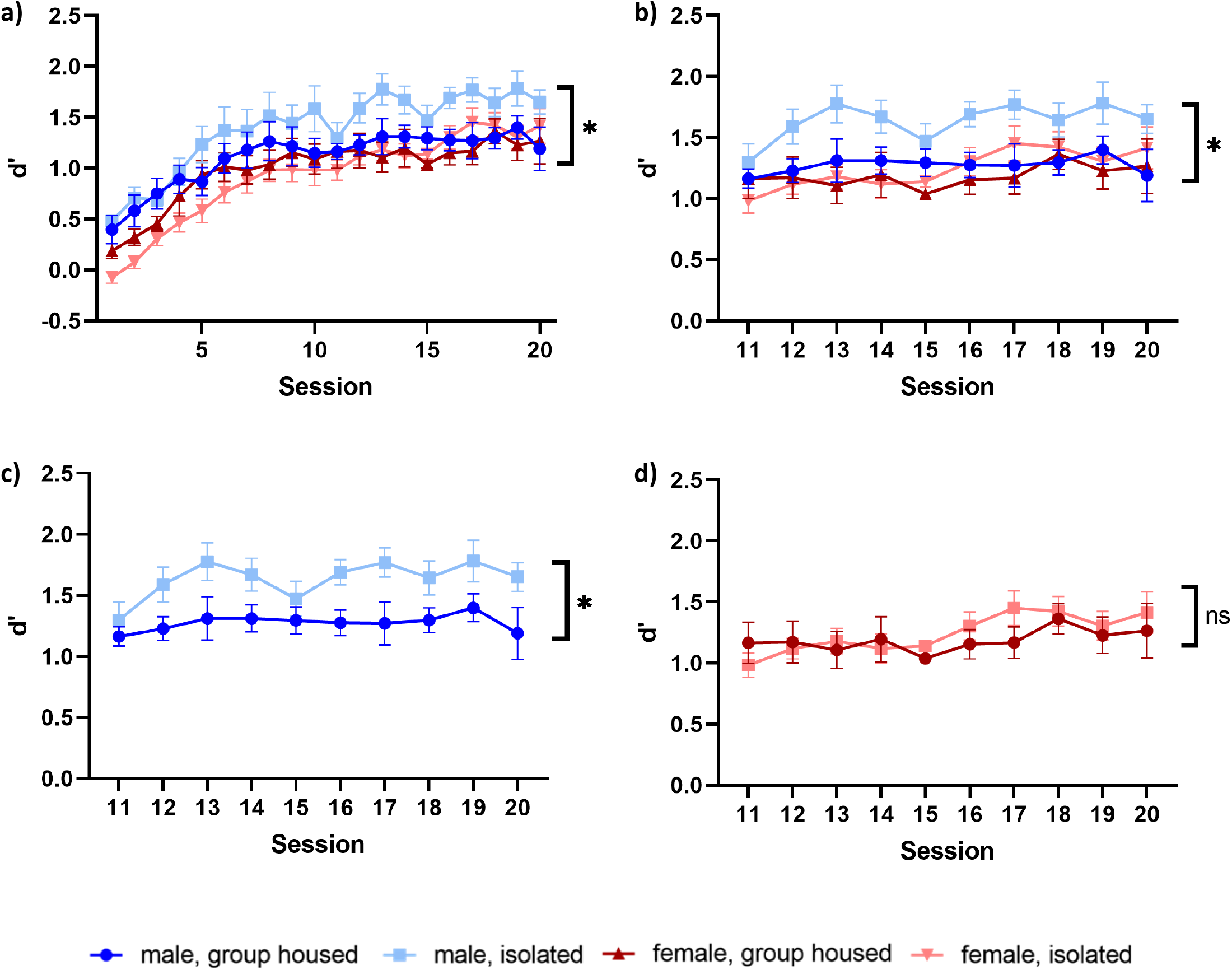
Performance in the continuous performance test is affected by SI (P21-35) in males. a) Discrimination index (d’) over the twenty sessions of stage 4 of CPT of the four groups. b) Discrimination index over the last seven sessions of stage 4 of CPT of the four groups. c) SI did not affect d’ in females. d) SI increased d’ in males. n = 6 GH males, 7 SI males, 6 GH females, and 8 SI females. Data are mean ± SEM. *p < 0.05

### 4.3 SI did not affect reaction time in males or females

Reaction time (RT) was measured between S+ presentation and screen touch (correct reaction time), S-presentation and screen touch (mistake reaction time), and screen touch and reward collection. Neither sex nor housing significantly affected correct RT (Figure 3a, Housing: F_1,23_ = 0.1155, p = 0.7371; Sex: F_1,23_ = 2.440, p = 0.1319). Mistake RT did not show a housing effect (F_1,23_ = 0.2791, p = 0.6023), but did show a sex effect, with females being slower to commit a mistake response than males (Figure 3b, F_1,23_ = 5.915, p = 0.0232). Reward collection time like correct RT was not affected by housing (F_1,23_ = 1.801, p = 0.1927) or sex (Figure 3c, F_1,23_ = 4.277, p = 0.0501).

**Figure 3.**
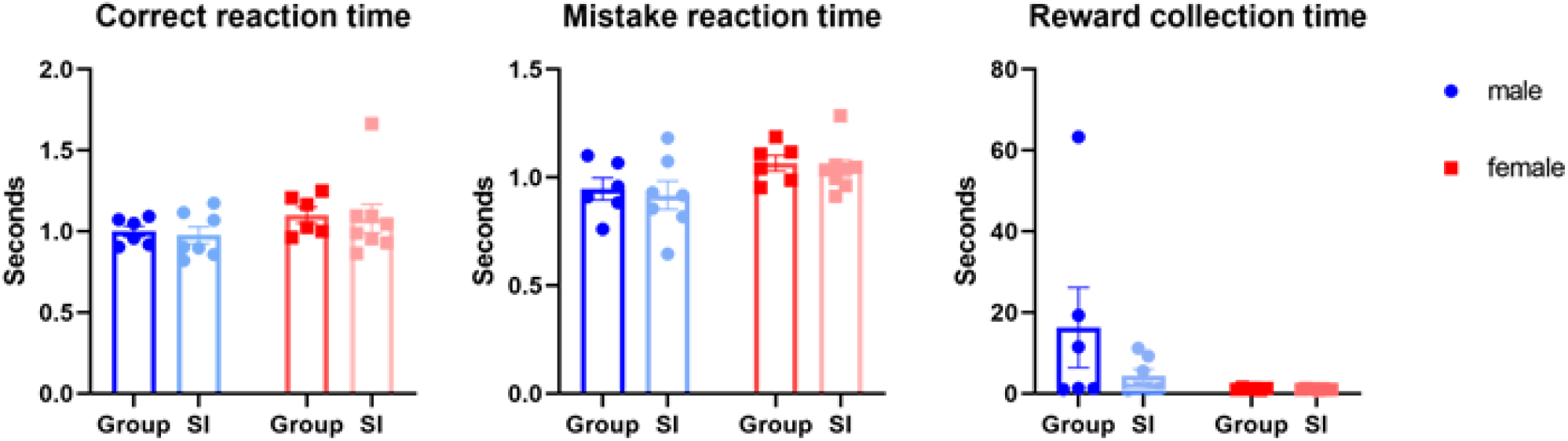
Reaction time (RT) in the continuous performance test is unaffected by sex or rearing. a) RT to S+ (hit) in stage 4 of CPT. b) RT to S- (mistake) in stage 4 of CPT. c) RT to reward collection after responding to S+ in stage 4 of CPT. n = 6 GH males, 7 SI males, 6 GH females, and 8 SI females. Data are mean ± SEM. *p < 0.05

### 4.4 Time bin analysis of CPT

CPT data was analyzed in three 15-minute bins (0-15, 15-30, and 30-45 minutes) to detect any changes in task performance over the duration of a session. (Figure 4) We found that performance was consistent among time bins for each housing condition, with SI males still scoring significantly higher than the three other groups. This effect reached significance in the first (Figure 4a, F_3,23_ = 4.458, p = 0.0131), second (Figure 4b, F_3,23_ = 3.971, p = 0.0204) and third (Figure 4c, F_3,23_ = 3.527, p = 0.0309) bins.

**Figure 4.**
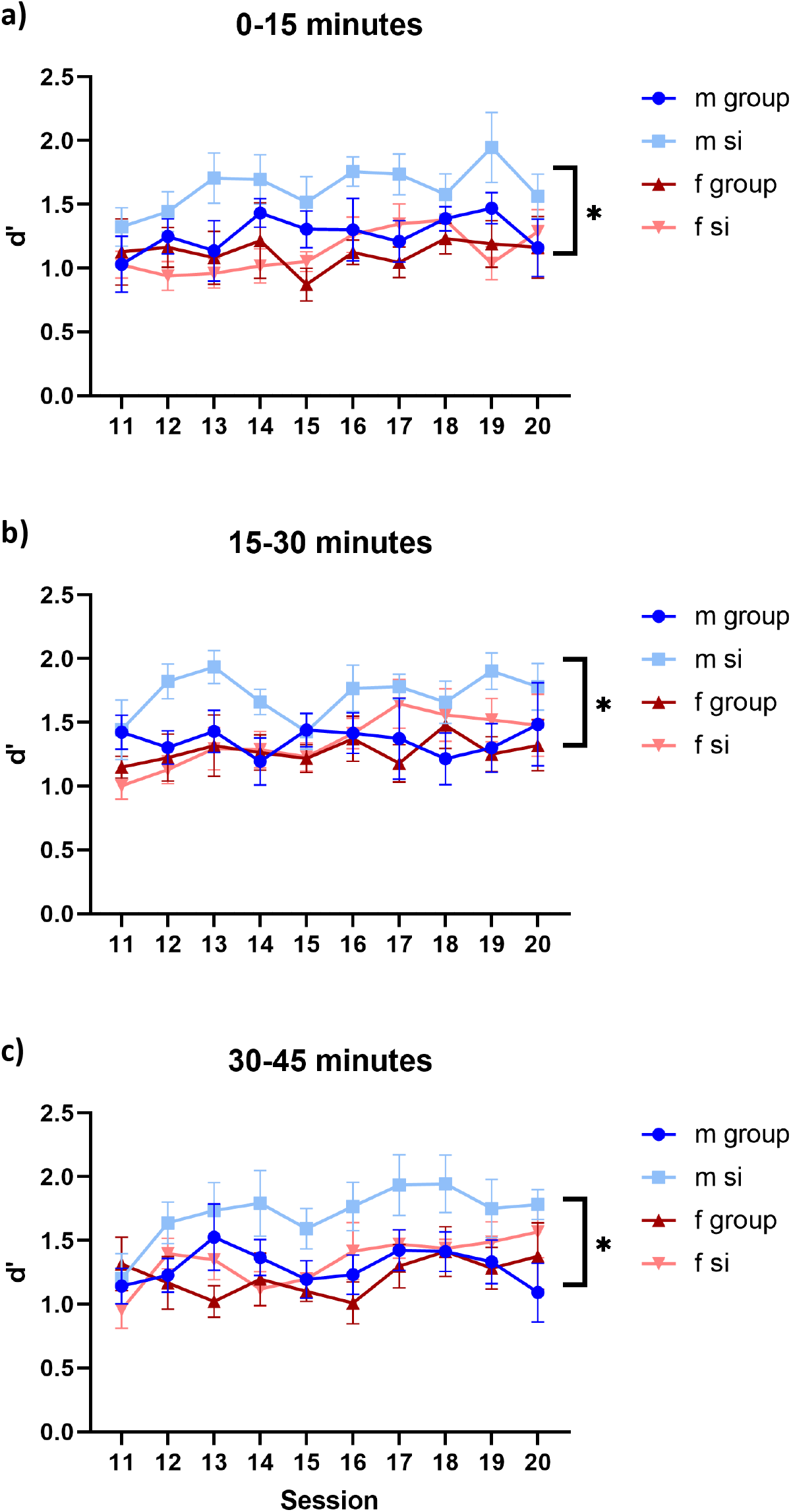
Time bin analysis of stage 4 of continuous performance test. a) 0-15 minutes. SI males scored significantly higher during the first 15 minutes of the CPT sessions. (p = 0.0084) b) 15-30 minutes. SI males scored significantly higher during the second 15-minute bin of CPT sessions. (p = 0.0722) c) SI males scored significantly higher during the last 15 minutes of the CPT sessions. (p = 0.0196) n = 6 GH males, 7 SI males, 6 GH females, and 8 SI females. Data are mean ± SEM. *p < 0.05

### 4.5 Male SI mice have higher break points in PR4 compared to group-housed controls

After observing that SI males scored higher in the CPT, a test sensitive to changes in attention, we tested males in a progressive ratio regimen, a battery sensitive to changes in motivation and reward-seeking behavior. This test allowed us to determine whether the observed CPT performance in SI males was being driven by an increase in attentional reserves or changes in reward-seeking behavior. We found that male mice that had undergone post-weaning SI expended more effort to obtain rewards during PR4 trials compared to group-housed littermates. (Figure 5a, F_13,11_ = 3.430, p = 0.0113)

**Figure 5.**
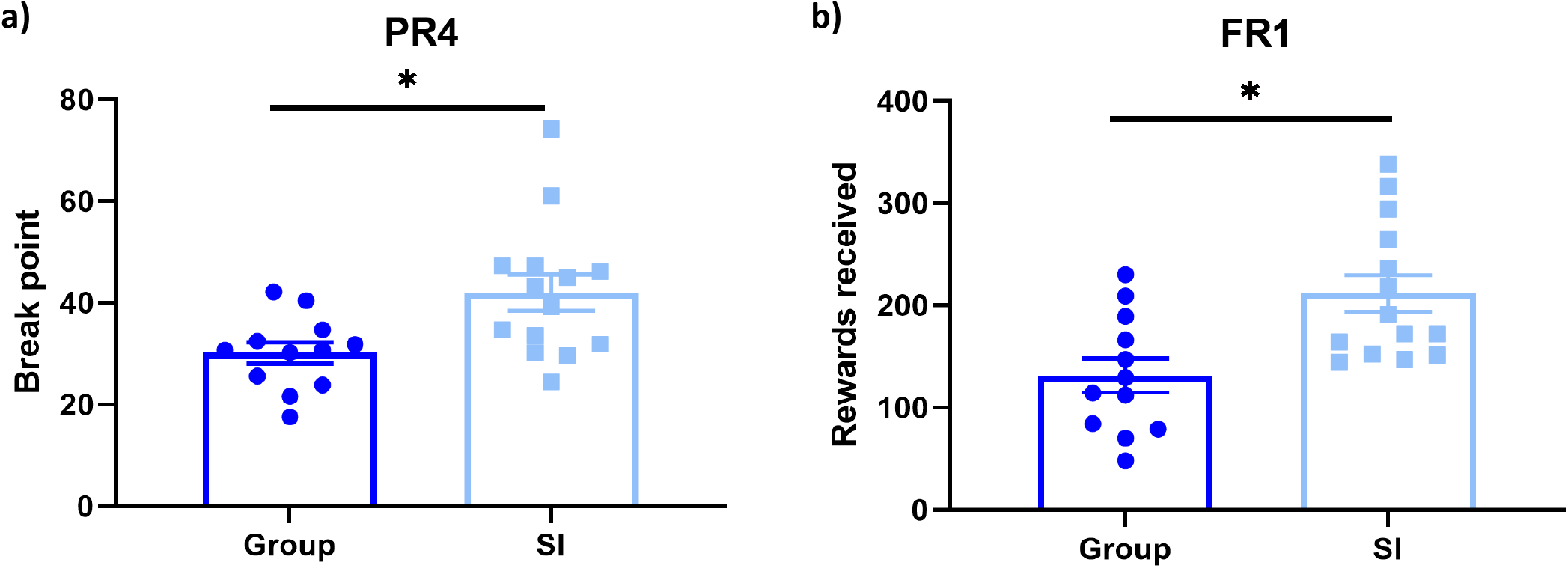
SI increases reward-seeking behavior in males as measured by a fixed/progressive ratio behavioral regimen. a) Average break point over seven PR4 sessions is increased in SI males. b) Reward intake during an unlimited FR1 session is increased in SI males. n = 12 GH males and 14 SI males. Data represent mean ± SEM. *p < 0.05

### 4.6 Male SI mice consume more reward in an uncapped FR1 session compared to group-housed controls

The motivation testing battery included an unlimited FR1 session between blocks of PR4 sessions to eliminate potential reward satiety as a confounding factor. We found that while both GH and SI mice consumed more rewards during the FR1 session than during the PR4 sessions, thus confirming that they were not reaching reward satiety, SI mice consumed significantly more rewards than their group-housed littermates. (Figure 5b, F_13,11_ = 1.349, p = 0.0037) The number of rewards received during this session also exceeded the amount of rewards received during any one session of the CPT, indicating that satiety was not a confounding factor in that test either.

## 5. Discussion

Adolescent social isolation causes measurable behavioral changes that persist through adulthood. Using a CPT, we found that SI increased sensitivity measures, usually interpreted as improved sustained attention, in male mice. SI also caused increases in reward-seeking behavior in males as measured by a FR and PR tasks, indicating the effects in the CPT may not have been specific to the domain of sustained attention. The effects of SI appeared to be sex-specific, as the performance of female mice in the CPT was unaffected by SI. These results together form a consistent, robust profile of the effects of adolescent SI on reward-seeking behavior.

### 5.1 Adolescent SI improves attention in males

This study investigated the effects of adolescent SI on sustained attention via CPT, which has established usage in rodent studies and human diagnostics [25]. Adolescent SI male mice performed significantly better on stage 4 of the CPT than group-housed males and both group-housed and SI females. Group-housed and SI females did not score significantly differently on the CPT.

To further characterize how mice engaged with the task, we analyzed the CPT session data using 15-minute time bins. Performance throughout each CPT session was consistent among the housing conditions, with male SI mice scoring significantly higher than the other housing conditions in all three bins. Female SI and group-housed mice again were not significantly different from each other. Higher scores in CPT are usually associated with improvements in sustained attention [23], but many studies of SI indicate impaired attention as a key feature of the phenotype [26]. In response to this finding, we considered performance in the CPT as an interaction between the mouse’s willingness to engage in the task and their interest in the food reward. If SI impaired attention but also increased reward-seeking behavior, this may read as an increase in CPT score depending on the weight of each change.

### 5.2 Adolescent SI increases reward-seeking behavior in males

To further investigate this interaction between attention and motivation, we utilized a fixed-ratio/progressive-ratio regimen. Only males were tested due to the fact that females did not show a significant effect of isolation in the CPT. Progressive ratio tasks measure effort committed to obtain a reward without the cognitively demanding rules of the CPT, so the PR results would possibly allow us to further refine the potential implications of the CPT results.

SI males showed increased effort in the PR, which is interpreted as increased reward-seeking behavior. This validates our hypothesis that changes in CPT performance caused by SI are due at least in part to significant differences in reward-seeking and motivation in SI males compared to the other groups tested. This finding is in agreement with the known role of isolation as a risk factor for substance use disorders in humans. [19]

### 5.3 Adolescent SI differentially affects males and females

The apparent resilience to SI stress in females was another finding of interest. Several other studies have found sex-specific responses to early life stress [27,28], including reduced spontaneous postsynaptic currents [27] and frontolimbic hyperconnectivity [28] that were observed in males and not in females. However, it is also possible that the cognitive tests chosen did not interrogate domains affected by SI in females.

### 5.4 Behavior effects in the greater context of adolescent SI effects

One of the largest remaining questions about the SI phenotype is how an organism translates the experience of SI stress to the observed behavioral effects. A previous study from our group [15] focused on the molecular effects of adolescent SI and found that the transcription factor ΔFosB was upregulated in SI males. Those findings about ΔFosB combined with the behavioral effects in this study are new pieces of the potential biological pathway between SI stress and the observed changes. Future experiments will directly test the causal relationship between SI, ΔFosB, and reward-seeking behavior. Increased reward-seeking behavior has also been observed following overexpression of ΔFosB. [17,29]

Our findings that transient isolation during adolescence affect males and females differently may provide critical information as we face the fallout from COVID-19-related SI. Although almost all people experienced increased isolation due to the pandemic, we may see significant variability in downstream effects Our data suggest that the long-term effects may not be immediately observable or evenly distributed among the population, but long-lasting and significant nevertheless.

## 6. Conclusions

This study tested the effects of postweaning SI on CPT and FR/PR performance in mice. SI exerts significant and sex-specific effects on performance in the tasks utilized in this study with male SI mice demonstrating increased reward-seeking and motivation for the palatable food reinforce across all tasks. These results are in line with previous studies on the effects of SI across species. Future studies will address the potential molecular and circuit intermediates that connect SI to behavioral changes.

## Acknowledgements

The authors thank Noelle White for technical assistance.

